# Gap junction coding innexin in *Lymnaea stagnalis*: sequence analysis and characterization in tissues and the central nervous system

**DOI:** 10.1101/785451

**Authors:** Brittany A. Mersman, Sonia N. Jolly, Zhenguo Lin, Fenglian Xu

## Abstract

Connections between neurons called synapses are the key components underlying all nervous system functions of animals and humans. However, important genetic information on the formation and plasticity of one type, the electrical (gap junction-mediated) synapse, is severely understudied, especially in invertebrates. In the present study, we set forth to identify and characterize the gap junction-encoding gene innexin in the central nervous system (CNS) of the mollusc pond snail *Lymnaea stagnalis* (*L. stagnalis*). With PCR, 3’ and 5’ RACE, and BLAST searches, we identified eight innexin genes in the *L. stagnalis* nervous system named *Lst Inx1-8*. Phylogenetic analysis revealed that the *L. stagnalis* innexin genes originated from a single copy in the common ancestor of molluscan species by multiple gene duplication events and have been maintained in *L. stagnalis* since they were generated. The paralogous innexin genes demonstrate distinct expression patterns among tissues. In addition, one paralog, *Lst Inx1*, exhibits heterogeneity in cells and ganglia, suggesting the occurrence of functional diversification after gene duplication. These results introduce possibilities to study an intriguing potential relationship between innexin paralog expression and cell-specific functional outputs such as heterogenic ability to form channels and exhibit synapse plasticity. The *L. stagnalis* CNS contains large neurons and a functionally defined network for behaviors; with the introduction of *L. stagnalis* in the gap junction field, we are providing novel opportunities to combine genetic research with direct investigation of functional outcomes at the cellular, synaptic, and behavioral levels.

**Summary Statement:** By characterizing the gap junction gene innexin in *Lymnaea stagnalis*, we open opportunities for novel studies on the regulation, plasticity, and evolutionary function of electrical synapses throughout the animal kingdom.

## Introduction

From simple reflexes to high cognitive functions including learning and memory, all nervous system operations rely on two main forms of synaptic communication to efficiently transmit signals: chemical (transmitter-mediated) and electrical (gap junction-mediated) (Ovsepian, 2017). Gap junctions form intercellular channels, providing direct and efficient means of communication by allowing quick movement of ions and small molecules (<1kd) between the cytosol of coupled cells (Qu and Dahl, 2002). Gap junctions have been identified in both vertebrates (named connexins) and invertebrates (named innexins: **in**vertebrate analog of con**nexins**) and exhibit structural homology despite their lack of sequence similarity. Originally thought to only have importance in invertebrates (Nagy et al., 2018), gap junctions are now known to be expressed throughout the mammalian nervous system and in various organs such as the heart, skin, kidney, eye, and inner ear (Dere and Zlomuzica, 2012). Mutations of gap junction-forming genes and associated proteins or dysfunction of gap junction activity are associated with many human diseases including cancer, deafness, oculodentodigital dysplasia as well as fear-related behaviors and learning and memory deficiencies (Abrams and Scherer, 2012; Bissiere et al., 2011; Dere and Zlomuzica, 2012; Mas et al., 2004).

The discovery of direct intercellular communication was first made in the invertebrate crayfish by Furshpan and Potter (Furshpan and Potter, 1957; Furshpan and Potter, 1959) and then in lobster (Watanabe, 1958). Structural evidence of the existence of a “gap”-like nexus near plasma membranes of adjacent cells was revealed with electron microscopy studies in various tissues and cells in both vertebrates and invertebrates (Dewey and Barr, 1962; Dewey and Barr, 1964; Farquhar and Palade, 1963), and molecular cloning and characterization of gap junction-forming genes were first made from tissues of human and rat (Kumar and Gilula, 1986; Paul, 1986). The presence of innexin has been established in all invertebrates except for sponges and echinoderms (Skerrett and Williams, 2017; Watanabe, 1958); however, extensive studies of gap junction-forming innexins have been severely restricted to a select few invertebrate model organisms, such as the fruit fly *Drosophila melanogaster*, the nematode *Caenorhabditis elegans*, and the medicinal leech *Hirudo verbana* (Beyer and Berthoud, 2018). Such restrictions limit the extent to which evolutionary and functional analyses can be made; full characterization of a gap junction system in a novel and easy-to-study invertebrate species is long overdue.

In this study, we introduce the freshwater pond snail, *Lymnaea stagnalis* (*L. stagnalis*), to the gap junction field. Like its sea slug counterpart, *Aplysia californica*, *L. stagnalis* belongs to the phylum Mollusca and class Gastropoda. Mollusca is the second-largest phylum of invertebrate animals, and many molluscs, such as the gastropod *A. californica*, the cephalopod *Octopus vulgaris*, and the cephalopod squid *Loligo pealeii*, have proven to be valuable resources and models for making significant fundamental neurobiological discoveries (Brunelli et al., 1976; Tasaki and Takenaka, 1963; Tricarico et al., 2014). *L. stagnalis* has been used in studies ranging from simple locomotive behaviors (Syed and Winlow, 1991) to highly complex processes like synaptogenesis (Dmetrichuk et al., 2006) and learning and memory (Lukowiak et al., 2003; Straub et al., 2006). Previously a widely understudied species, recent efforts have established a transcriptome (Feng et al., 2009; Sadamoto et al., 2012) and genome (Davison et al., 2016) assembly of *L. stagnalis*, making molecular and genetic research of the organism even more applicable and providing an invaluable tool for future work in the field of molecular neurobiology.

The *L. stagnalis* central nervous system (CNS) has been well described, and established neuronal networks are available including morphological features, spatial topology, and types of synaptic connections (van Nierop et al., 2006). Importantly, *L. stagnalis* brain contains many gap junction-forming neurons that form a well-defined network for various behaviors. For example, the pedal dorsal A cluster neurons in the left and right pedal ganglia form gap junctions (electrical synapses) that control the cilia of the foot for locomotion (Kyriakides et al., 1989), and the A cluster motoneurons in the left and right cerebral ganglia form gap junctions that control whole body withdrawal response (Syed and Winlow, 1991). While well-defined neuronal networks including axon projections, synapse formation, and functional outcomes are known, most studies and knowledge of gap junctions in *L. stagnalis* and other species in the Mollusca phylum are limited to electrophysiological and behavioral work with little genetic information (Carrow and Levitan, 1989; Dargaei et al., 2014; Vehovszky and Elliott, 2000).

For the first time in *L. stagnalis*, we identified eight innexin genes named *Lst Inx1-8* (accession numbers are provided in Table 1). Phylogenetic analyses revealed the origin and evolutionary history of the eight paralogs in Mollusca. The expression pattern of one innexin, *Lst Inx1*, was analyzed via *in situ* hybridization (ISH) and demonstrated variable localization within ganglia that contain single cells known to form electrical synapses. Such information provides a necessary foundation for future investigation of the genetic and molecular mechanisms of nervous system development and function in *L. stagnalis* and other invertebrate species.

**Table 1.** Accession numbers of innexin genes in *L. stagnalis*

| Gene Name | Accession Number |
| --- | --- |
| *Lst Inx1* | MN480796 |
| *Lst Inx2* | MN480797 |
| *Lst Inx3* | MN480798 |
| *Lst Inx4* | MN480799 |
| *Lst Inx5* | MN480800 |
| *Lst Inx6* | MN480801 |
| *Lst Inx7* | MN480802 |
| *Lst Inx8* | MN480803 |

## Materials and Methods

### Animals and CNS dissection

The freshwater snails, *Lymnaea stagnalis*, were kept in artificial pond water at 20-22°C on a 12 h light/dark regimen and fed romaine lettuce. Snails ∼12 months old were used for innexin sequence identification, tissue expression, and ISH experiments. CNS isolation was performed as previously described (Syed et al., 1990). Briefly, snails were de-shelled and anaesthetized in Listerine solution (21.9% ethanol and 0.042% methanol; department store; everywhere) diluted to 10% in *Lymnaea* saline (51.3 mM NaCl; 1.7 mM KCl; 4.0 mM CaCl2; 1.5 mM MgCl2, 10 mM HEPES, pH 7.9). Dissected central ring ganglia was used for either RNA or genomic DNA (gDNA) extraction.

### RNA and gDNA extraction

RNA was extracted from the central ring ganglia and tissues of *L. stagnalis* with the RNeasy Mini Kit (Qiagen; 74104; Venlo, Netherlands) according to the manufacturer’s instructions. An additional DNase digestion step (Qiagen; 79254; Venlo, Netherlands) was added to the protocol to prevent DNA contamination. gDNA was used for normalization. gDNA extraction (Invitrogen; K1820-02; Carlsbad, CA, USA) was completed according to the manufacturer’s instructions.

### Identification of innexin genes in *L. stagnalis*

To determine whether homologs of innexin are present and expressed in *L. stagnalis*, we reverse transcribed RNA extracted from whole CNS to cDNA with Superscript II Reverse Transcriptase (Invitrogen; 18064-014; Carlsbad, CA, USA). PCR was then performed with an Eppendorf Mastercycler Gradient 5331 (Hauppauge, NY, USA), Taq DNA polymerase (New England Biolabs; M0273A; Ipswich, MA, USA), and degenerate primers designed for innexin detection in the crab *Cancer borealis* (Shruti et al., 2014): forward primer 5’-GAGGACGAGATCAA-GTACCACACATAYTAYCARTGG-3’ and reverse primer 5’-GGCATGAAGGTCAGGAA-GACGWRCCARAACC-3’. Because innexin genes contain regions rich in sequence conservation among all invertebrates (Beyer and Berthoud, 2018), it is interesting, but not surprising, that the degenerate primers designed for amplification in *C. borealis* also amplified a partial sequence in *L. stagnalis* (Fig. S1). The partial fragment was sequenced (Genewiz; South Plainfield, NJ, USA) and used to design primers (Table 2) for 3’ (Invitrogen; 18373-027; Carlsbad, CA, USA) and 5’ (Invitrogen; 18374-041; Carlsbad, CA, USA) rapid amplification of cDNA ends (RACE) to obtain a complete mRNA transcript. We named the gene *Lst Inx1* (Table 1) according to common innexin naming strategies. We then used the translated amino acid sequence of *Lst Inx1* as a query in NCBI BLAST to search for its orthologous genes in other species. The top hits of the BLAST search were *Aplysia californica pannexin1, Biomphalaria glabrate innexin unc-9 like*, and *Crassostrea gigas innexin unc-9*, which further supported that *Lst Inx1* belongs to the innexin gene family.

**Table 2.**
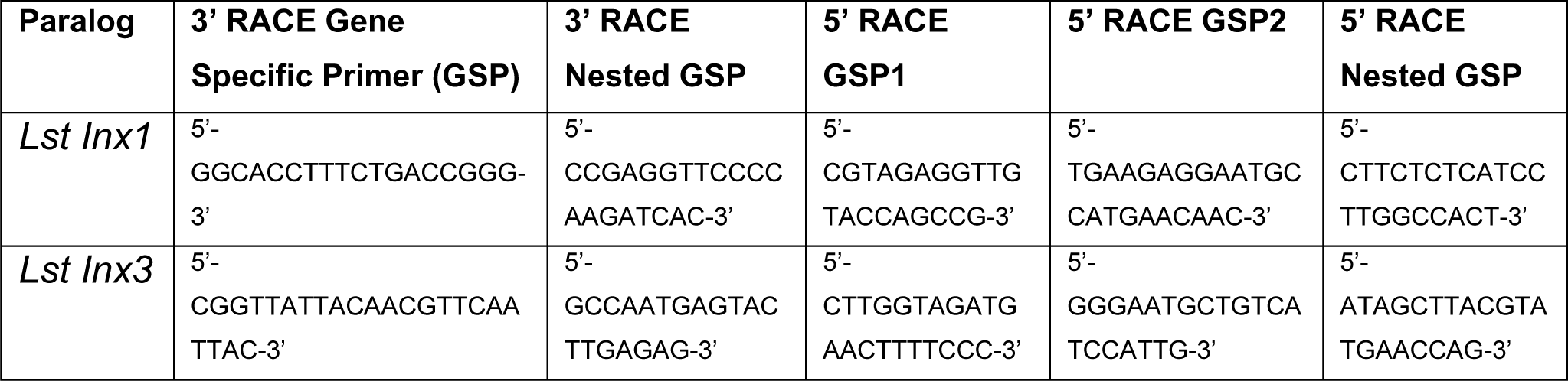
Primers used for 3’ and 5’ RACE

To determine whether paralogous genes of *Lst Inx1* are present in the *L. stagnalis* genome, we then used the *Lst Inx1* amino acid sequence as a query to run a TBLASTN search against the genome sequence of *L. stagnalis* (assembly v1.0) from the NCBI WGS database (Skerrett and Williams, 2017). The BLAST search identified 10 significant hits (E value < 1e-10, alignment region > 50% of query). Because the genome assembly of *L. stagnalis* is highly fragmented (328,378 scaffolds with N50=5751), to determine whether each hit represented a unique genomic locus, we examined the genomic context for each hit. Three hits were found in three scaffolds that share 99.9% of sequence identities. The three hits were thus considered the same gene. Therefore, we identified eight paralogous genes of innexin in *L. stagnalis*, aptly named *Lst Inx1* through *Lst Inx8*. The nucleotide sequences of these innexin genes were translated to amino acid sequences via ExPASy translation tool. 3’ and 5’ RACE (Table 2) was completed as previously described on *Lst Inx1*, and a complete ORF was obtained. The 3’ and 5’ ends of the remaining six genes were predicted via homology studies utilizing other invertebrate species’ innexin sequences including *B. glabrate* and *A. californica*. Three of the predicted genes, *Lst Inx2*, *Lst Inx5*, and *Lst Inx6*, were validated via PCR and primers designed in the first and last exon of each predicted sequence (Table 3). The eight sequences were then used in a multiple sequence alignment generated by T-Coffee (Fig. 1), and a second multiple sequence alignment was generated with *Lst Inx1* and an innexin ortholog in *C. elegans* and *D. melanogaster*, CELE R07D5.1 and Dmel CG4590 INX2, respectively (Fig. 2A) (Di Tommaso et al., 2011; Notredame et al., 2000).

**Figure 1.**
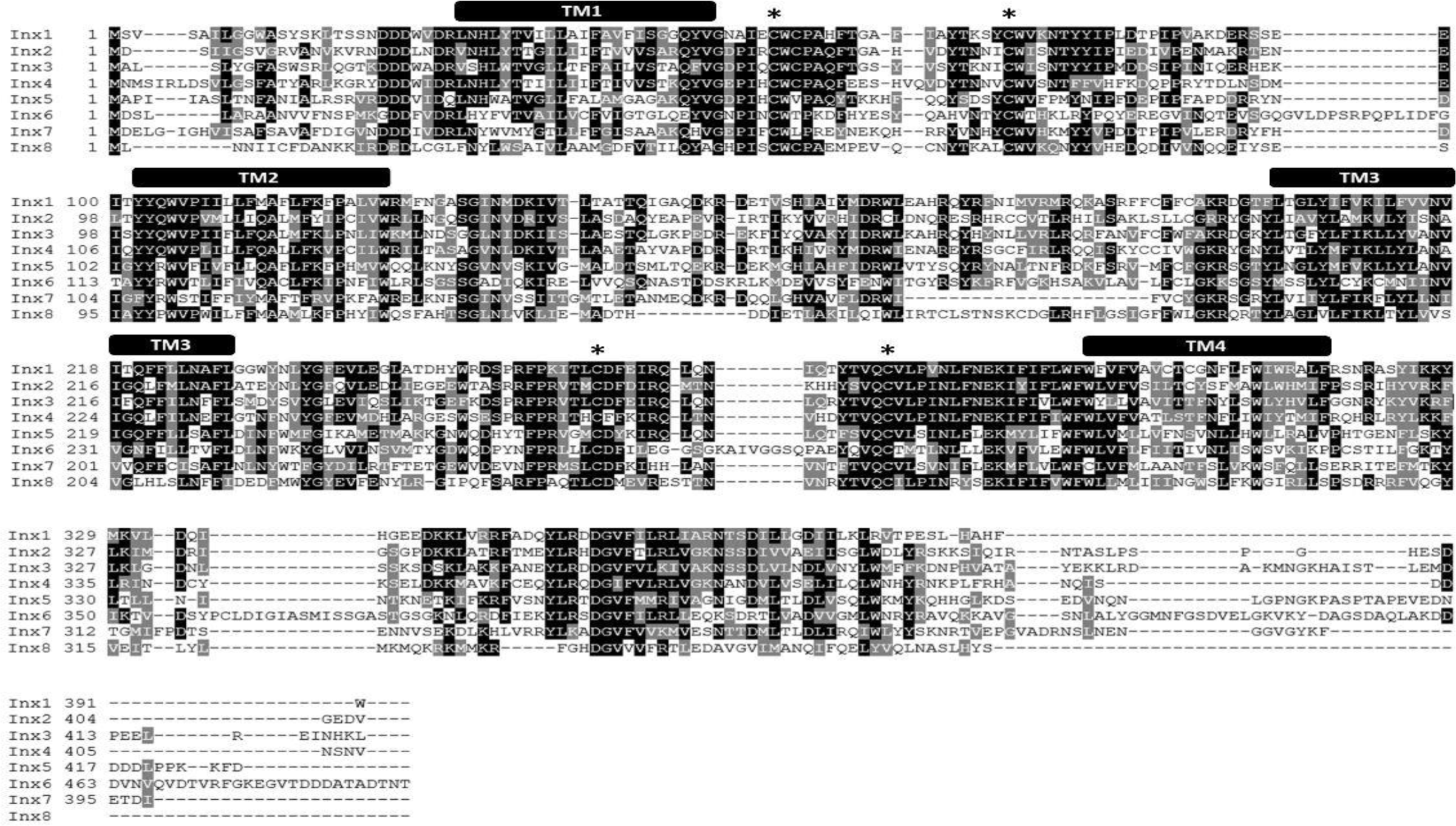
*L. stagnalis* express multiple paralogs of innexin in their CNS. RACE and PCR experiments revealed eight innexin paralogs within the *L. stagnalis* genome. Amino acid alignment revealed conserved residues among the paralogs. Transmembrane domains are indicated above the sequences, and asterisks (*) indicate the two cysteines conserved across all innexins located in the two extracellular loops.

**Figure 2.**
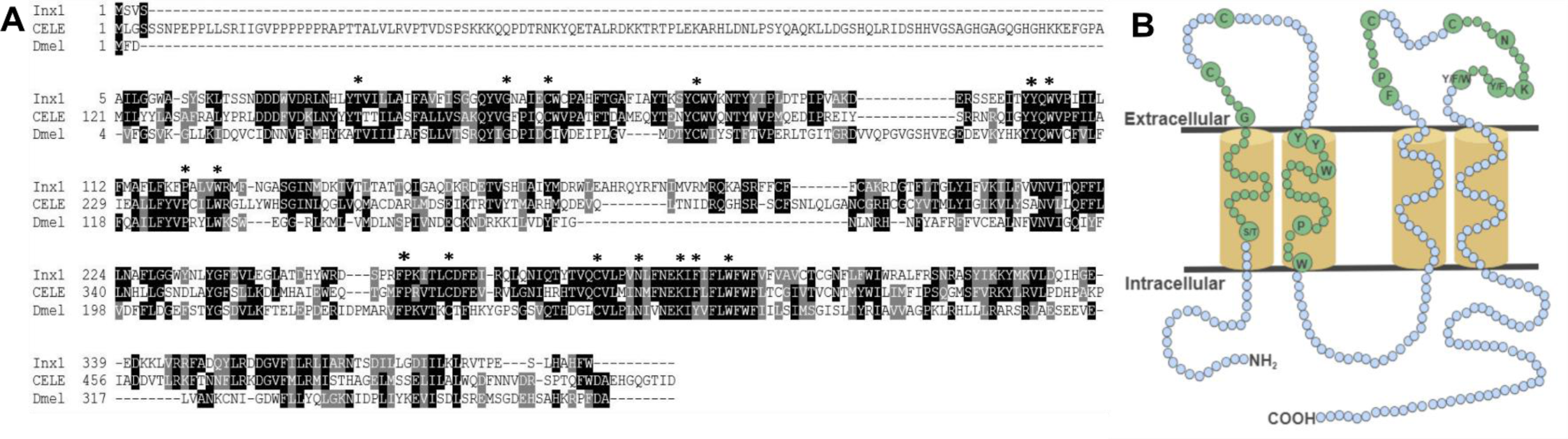
The conserved topology and sequences of innexins. **A.** A multiple sequence alignment of *Lst Inx1* with one innexin in *C. elegans* and *D. melanogaster* shows the amino acid residues conserved in all innexins (shown in B) **B.** In this model, cylinders are transmembrane domains, and circles represent amino acids. Small, blue circles signify a variation in number of amino acids while small, green circles signify an invariable number of amino acids. Residues conserved across all innexin sequences are written in big, green circles and are highlighted by an asterisk (*) in A.

**Table 3.**
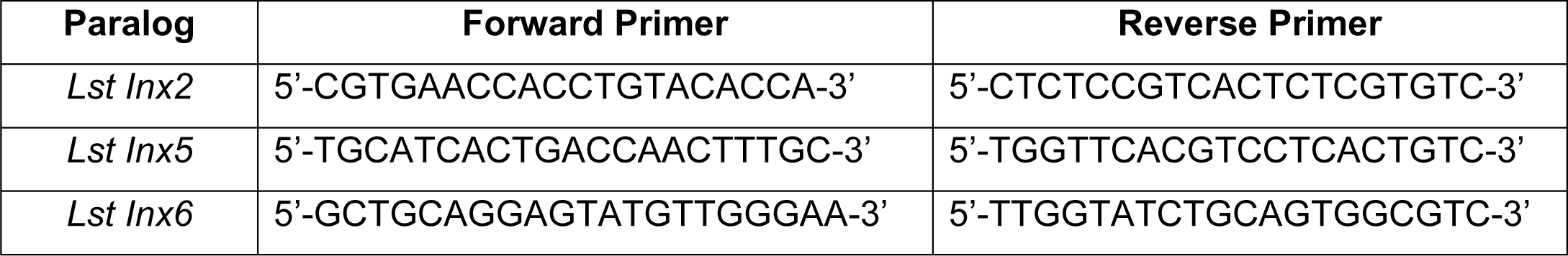
Primers used for validation of predicted paralog sequences

### Phylogenetic analysis

We used the amino acid sequence of *Lst Inx1* as a query to search for its homologous sequences via NCBI BLASTP in representative invertebrate species: the Molluscs *Aplysia californica*, *Pomacea canaliculata*, and *Octopus bimaculoides*; the Annelida species *Helobdella robusta*; the Arthropod *D. melanogaster*; and the Nematode *C. elegans*. A total of 89 innexin homologous genes were obtained from the seven species (E value < 1e-5, alignment region > 50% of the query). The phylogenetic tree of the innexin gene family was inferred by using the maximum likelihood (ML) method implemented in MEGA 7 (Kumar et al., 2016). The best substitution model for innexin sequences was inferred by the ML fit test tool in MEGA 7 (LG+G+I, α = 1.46).

### Tissue expression analysis of innexins via reverse transcription (RT-PCR)

To identify expression of innexin paralogs across the body of *L. stagnalis*, RNA and gDNA was extracted from various tissues: CNS, buccal mass, penis, albumin gland, and foot. Seven ∼12-month-old snails held in the same aquatic tank were chosen at random for dissection of tissue. The CNS was dissected from all seven snails, and other tissues were dissected from five of the snails. One CNS and two buccal mass samples were excluded from the experiment due to poor quality RNA. RT-PCR was performed with Superscript III One-Step Platinum Taq (Invitrogen; 12574-026; Carlsbad, CA, USA) and primers designed to amplify each paralog (Table 4). All gDNA extracted from the tissues underwent the same reactions for normalization; amount of template, primers used, and Mastercycler conditions were kept constant for all RNA and gDNA reactions. A 1% agarose gel was used to determine innexin paralog expression. All gel images were taken with the ChemiDoc MP Imaging System (BioRad; 12003154; Hercules, CA, USA). Expected sizes of amplified products were *Lst Inx1* (498bp), *Lst Inx2* (170bp), *Lst Inx3* (404bp), *Lst Inx4* (316bp), *Lst Inx5* (147bp), *Lst Inx6* (164bp), *Lst Inx7* (446bp), and *Lst Inx8* (404bp). The intensity of expression (band intensity) was calculated with ImageJ software for both RNA and gDNA PCR products. The log2 ratio of RNA to gDNA band intensity was calculated to represent relative gene expression as previously described (Tsankov et al., 2010). A no-RT control was used by heating the Superscript III Platinum Taq mix at 95°C for 5 min to inactivate the enzyme according to the manufacturer’s instructions.

**Table 4.**
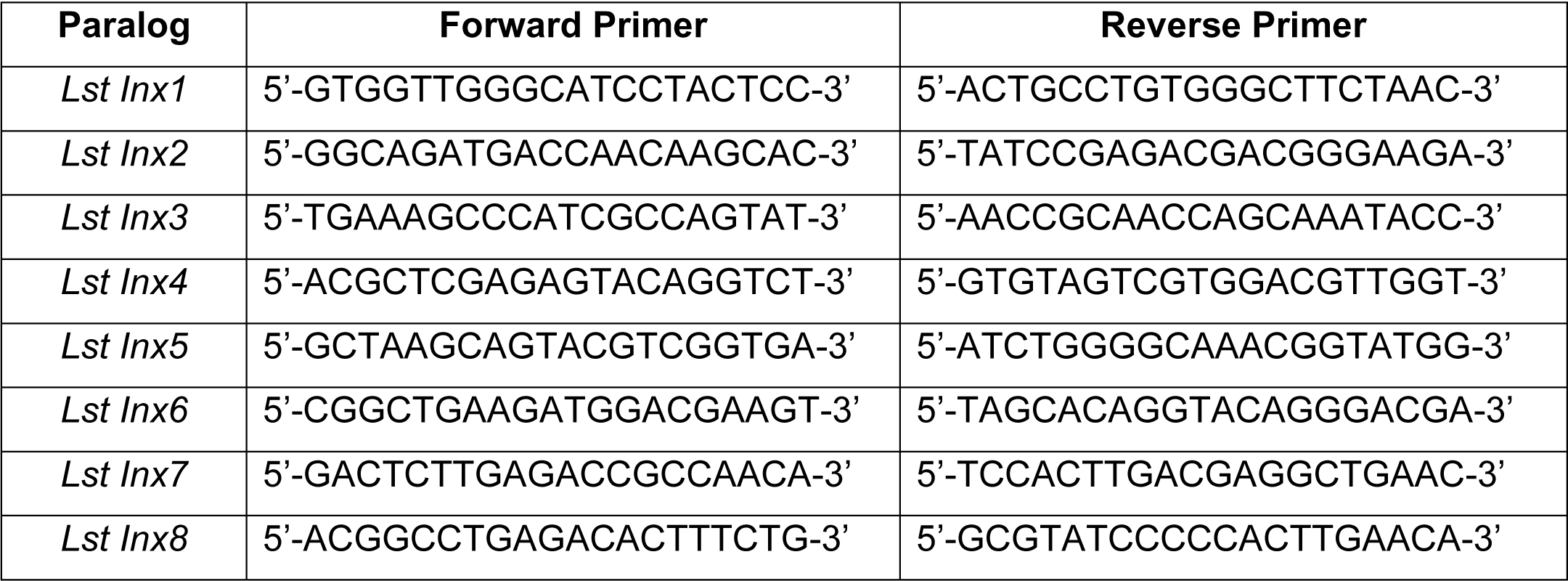
Primers used for tissue expression analysis

### Quantification of transcription abundance of innexin genes based on RNA-sequencing data

We downloaded the raw RNA-sequencing data of the CNS in *L. stagnalis* from the NCBI SRA database (SRA ID DRX001464). This dataset consists of 81.9 million single-end reads with a read length of 100 nucleotides. The RNA-seq reads were mapped to the genome sequence of *L. stagnalis* (assembly v1.0) using HISAT (Kim et al., 2015). 73.52% of reads were aligned to the *L. stagnalis* genome exactly one time. The numbers of reads mapped to each innexin gene were counted by using the “featureCounts” (Liao et al., 2014). The transcription abundance of each innexin gene was normalized as Reads Per Kilobase of transcript, per Million mapped reads (RPKM, Table S1).

### *In situ* hybridization

To assess the localization pattern of *Lst Inx1* throughout *L. stagnalis* CNS, ISH was performed with digoxigenin (DIG)-labeled probes. Twelve-month-old snails were randomly chosen and anesthetized, and the CNS was dissected. The commissure connecting the left and right cerebral ganglia was cut to allow the entire CNS to be splayed flat. Each CNS sample was paraffin embedded and sectioned into four ∼10µm slices. After sectioning, the samples were washed with xylene three times to dewax and rehydrated through an ethanol series (100% for two washes and 95%, 90%, 80%, 70%, diH2O for one wash each). To allow hybridization, samples were fixed with 4% paraformaldehyde for 20 min and washed twice with DEPC-PBS for 5 min each.

Samples were then treated with proteinase K (50µg mL^-1^) at 37°C for 13 min. Samples were again washed in DEPC-PBS, fixed with 4% paraformaldehyde, and rinsed with DEPC-H2O. Pre-hybridization solution (BioChain; K2191050; Newark, CA, USA) was added to the samples for 4 hours at 50°C followed by incubation in 4ng mL^-1^ DIG-labeled probe (Table 5) at 45°C overnight. Probes for *Lst Inx1* were designed to target regions with little sequence similarity between the eight paralogs, indicating the localization of *Lst Inx1* transcript alone. Two probes were used against *Lst Inx1*, and the experiment was repeated three times with four snails each experiment to ensure reliability; one probe targeted nucleotide 273 region while a second targeted nucleotide 601 region with sense probes acting as controls. Samples were washed with 2XSSC, 1.5XSSC, and 0.2XSSC and incubated with blocking solution for one hour at room temperature. To visualize the transcript, samples were incubated with AP-conjugated anti-DIG antibody for 4 hours, washed with PBS and alkaline phosphatase buffer, and incubated with NBT and BCIP in alkaline phosphatase buffer overnight. After rinsing with diH2O, phase contrast images were taken on an inverted microscope (Olympus CKX53; Bridgeport, CT, USA). Images are shown in Fig. 5.

**Table 5.**
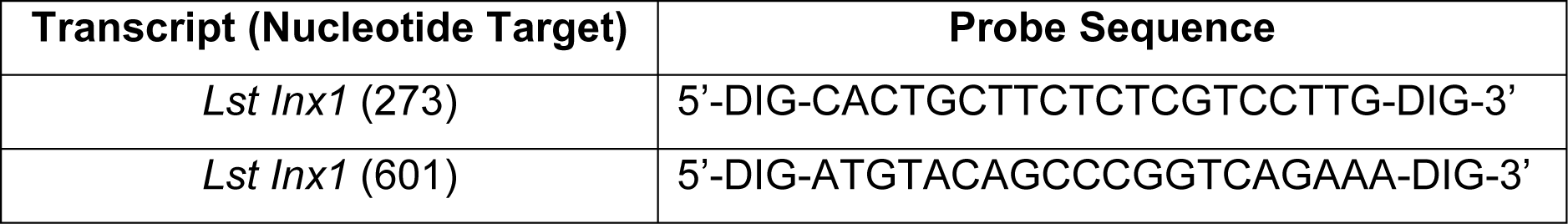
DIG-labeled probes used for *in situ* hybridization

## Results

### Sequence comparison of *L. stagnalis* innexin paralogs

RNA extracted from the CNS of *L. stagnalis* revealed eight paralogs of innexin, named *Lst Inx1* through *Lst Inx8*. The innexin sequences were transcribed and used to create a multiple sequence alignment with T-Coffee (Fig. 1). Comparison of trends in the alignment and commonly conserved amino acids in innexins (Fig. 2B) strengthened our confidence in the confirmed and predicted *L. stagnalis* sequences. For example, all invertebrate innexins share two strictly conserved cysteines in each extracellular loop and a YY(x)W region in the second transmembrane domain (Phelan and Starich, 2001); the eight innexin paralogs identified in *L. stagnalis* also shared these conserved regions. Topology studies with membrane-spanning protein prediction software revealed the expected four transmembrane structure of typical gap junction proteins (Beyer and Berthoud, 2018). A separate multiple sequence alignment (Fig. 2A) comparing *Lst Inx1* with the two most conserved innexin orthologs in *C. elegans* and *D. melanogaster*, CELE R07D5.1 and Dmel CG4590 INX2, respectively, revealed conserved amino acid residues common in innexins of all invertebrates (Fig. 2B).

### Phylogenetic analysis of the origin and evolution of innexin genes in *L. stagnalis*

To infer the origin and evolution of the eight innexin genes in *L. stagnalis*, we reconstructed a phylogenetic tree using the amino acid sequences from seven representative invertebrate species (Materials and Methods; Fig. 3A). The ML phylogenetic tree shows that all innexin genes from the four Mollusca species, including *L. stagnalis*, *A. californica*, *P. canaliculata*, and *O. bimaculoides*, form a well-supported monophyletic clade, suggesting a single origin of innexin genes in Mollusca. These Mollusca innexin genes form seven well-supported clades (Fig. 3, indicated by shaded regions), and each of the clades contain members from at least three Molluscan species. This topology suggests that multiple gene duplication events of innexin have occurred prior to the divergence of Mollusca, which generated at least five copies of innexin genes. One ancestral innexin gene was further duplicated before the divergence of *L. stagnalis*, *A. californica*, and *P. canaliculate* that generated *Lst Inx6-8*. Like Mollusca, innexins in other phyla, Annelida, Arthropoda, and Nematoda, also form a phylum-specific clade, suggesting that they originated from a single ancestral gene copy in each phylum and multiple gene duplication events followed (Fig. 3B). The similar expansion patterns in major invertebrate phyla suggests that duplication and functional diversification of innexin genes might have played an important role in phylum-specific electrical synapse function and nervous system development.

**Figure 3.**
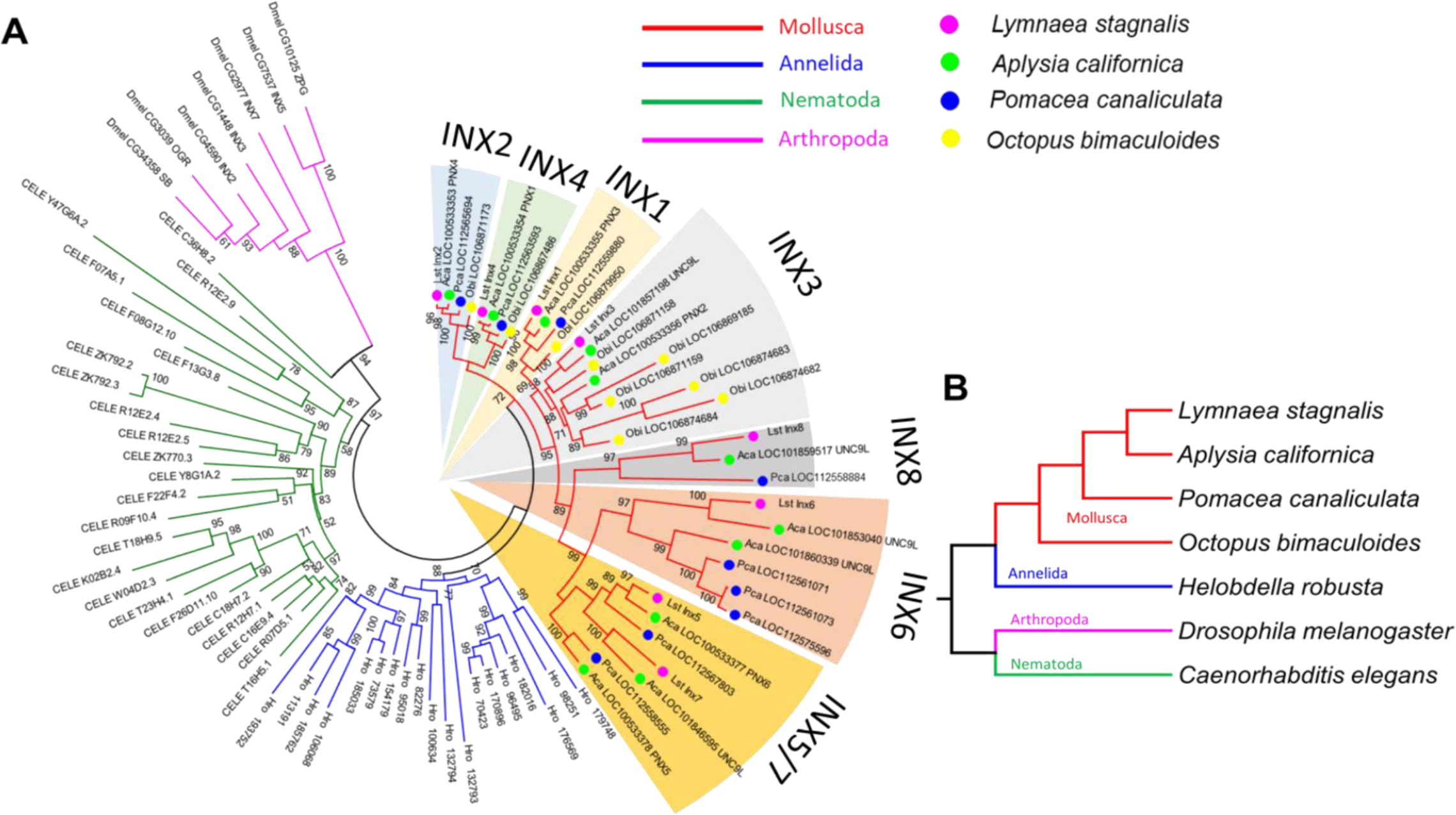
*L. stagnalis* innexins are evolutionarily related to innexins in other invertebrates. **A.** Phylogenetic analysis revealed the evolutionary relationship between *L. stagnalis* innexin paralogs and innexin orthologs in other invertebrates including species within the Mollusca family, to which *L. stagnalis* belongs, and well-studied species within Annelida, Nematoda, and Arthropoda. The different families are separated by branch color: Mollusca (red), Annelida (blue), Nematoda (green), and Arthropoda (pink). The Mollusca family is further sorted by colored circles: *L. stagnalis* (pink), *A. californica* (green), *P. canaliculata* (blue), and *O. bimaculoides* (yellow). Shading is used to indicate the seven well-supported clades formed in Mollusca. **B.** A phylogenetic tree demonstrates the evolutionary relationship between the species analyzed. The same branch color scheme is used as in A.

### Expression of innexin throughout *L. stagnalis* tissues

To gain a better understanding of the expression patterns of each innexin paralog throughout the body of *L. stagnalis*, we performed RT-PCR with primers specific to each paralog. Five organs of the snail were tested for specific reasons. The CNS was hypothesized to have very high levels of innexin expression. The buccal mass and foot are innervated by two sets of electrical synapse-forming cells, the octopamine neurons to regulate feeding and left/right pedal A neurons to regulate locomotion, respectively (Kyriakides et al., 1989; Vehovszky and Elliott, 2000). The albumin gland secretes epidermal growth factor required for synapse formation (Munno et al., 2000). The penis was used in a similar experiment testing the presence of nicotinic acetylcholine receptor expression (van Nierop et al., 2006) and was also used here. After RT-PCR with reactions using RNA or gDNA as starting material, agarose gel electrophoresis was completed (Fig. S2), with inactivated reverse transcriptase reactions used as controls.

Relative expression of each paralog was calculated throughout all tissues (Materials and Methods, Fig. 4), and a heatmap was created to demonstrate the changes in paralog expression between tissue type. Our results show that innexin genes are ubiquitously expressed throughout the entire body of *L. stagnalis*, and no obvious tissue-specific trends were imminent. However, innexin paralogs are up- and down-regulated throughout the same tissue. For example, in sample CNS-1, *Lst Inx2* is upregulated while *Lst Inx7* is downregulated. Interestingly, *Lst Inx7* had noticeably less expression throughout all tissues tested. *Lst Inx4* and *Lst Inx8* also had generally lower expression in all tissue types while *Lst Inx2* had higher expression throughout the tissues.

**Figure 4.**
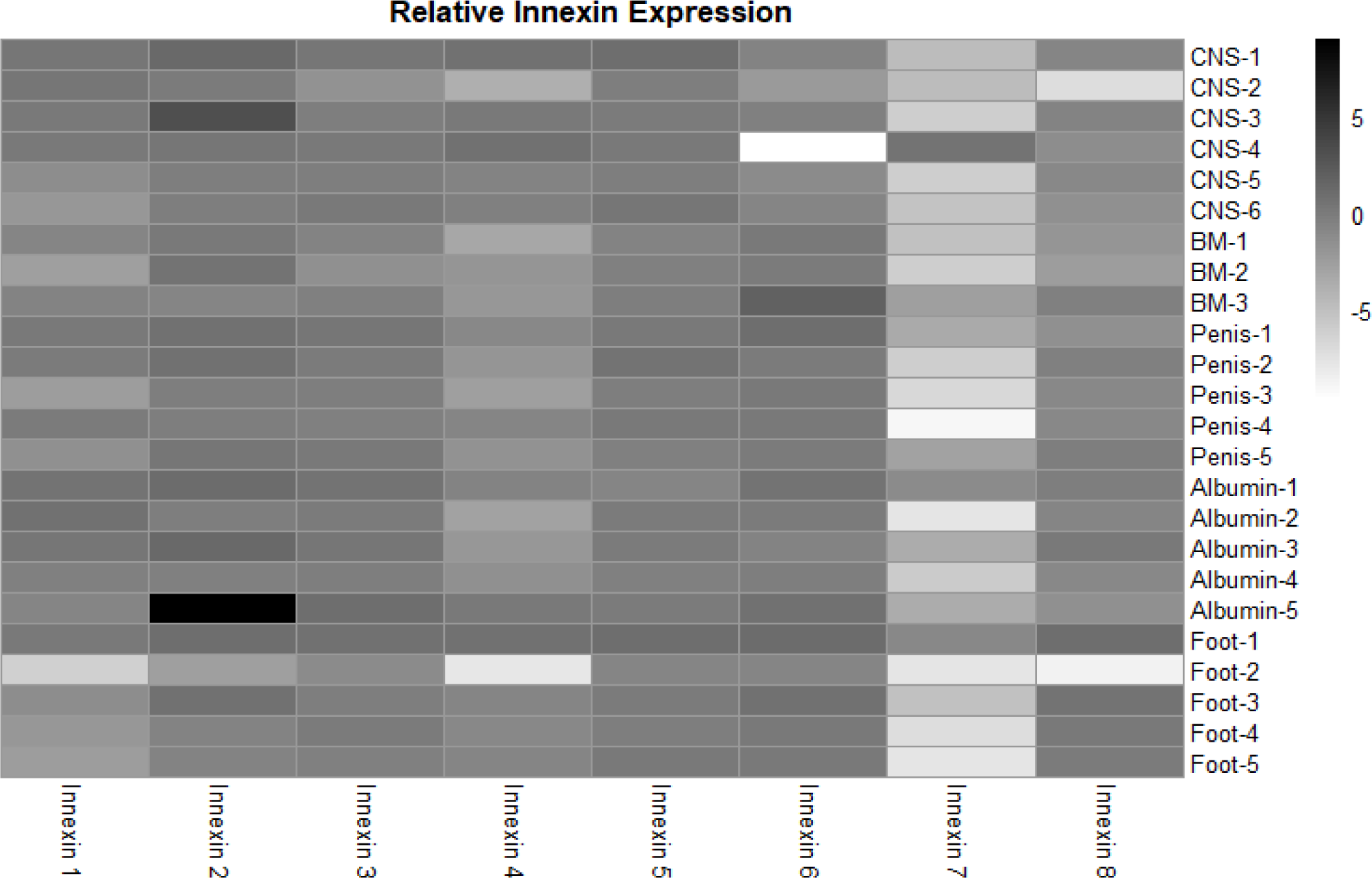
Analysis of relative innexin expression in different tissue types revealed both up- and down-regulation of innexin paralogs. RNA and gDNA from *L. stagnalis* was extracted from five tissue types and used in RT-PCR to determine relative innexin expression. Overall, the results showed variable expression between paralogs. One paralog, *Lst Inx7*, was down-regulated in most tissues with greater than 5-fold down-regulation in many cases. Sample sizes are as follows: CNS (n=6), buccal mass (n=3), penis (n=5), albumin gland (n=5), foot (n=5).

**Figure 5.**
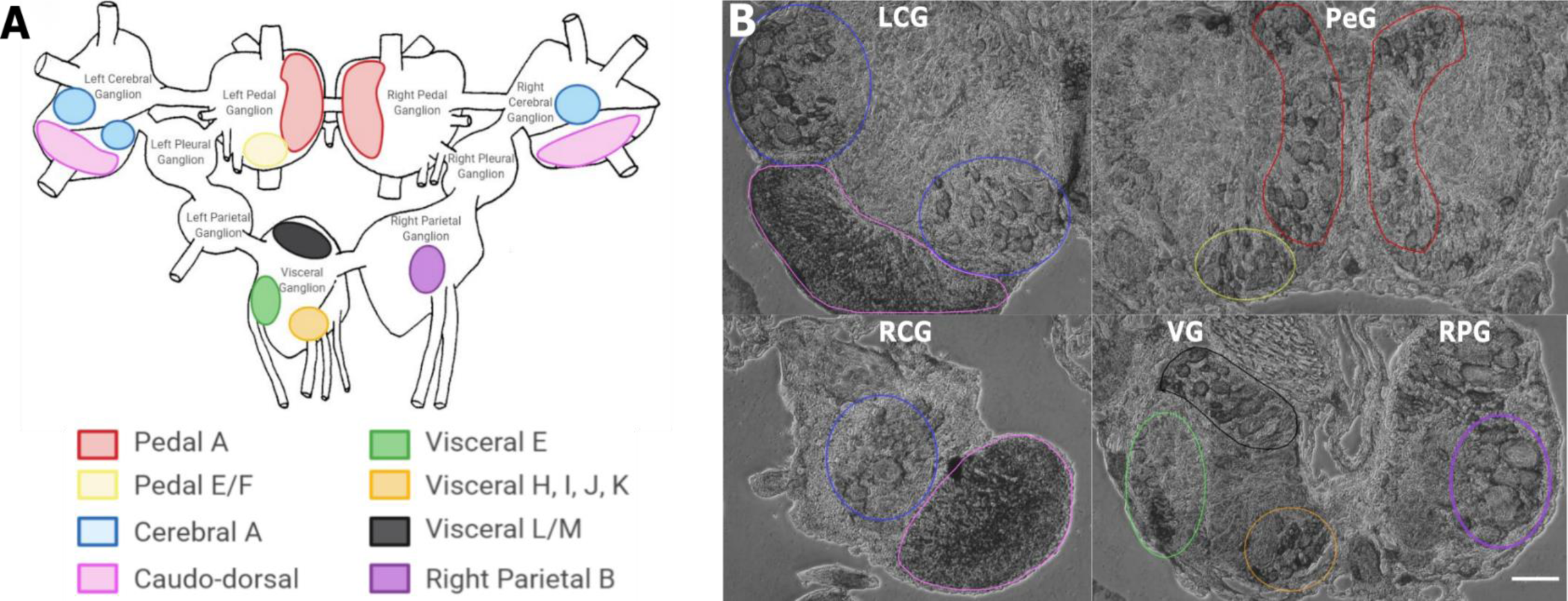
*In situ* hybridization with probes targeting *Lst Inx1* demonstrated localization of the transcript to neuron clusters throughout the *L. stagnalis* CNS. **A.** A schematic of the *L. stagnalis* CNS shows the location of the nine ganglia and highlights *Lst Inx1*-positive clusters. **B.** ISH revealed localization of *Lst Inx1* mRNA in regions of specific ganglia, with colors in A corresponding to the same colored outline of clusters in B. LCG: left cerebral ganglion, RCG: right cerebral ganglion, PeG: pedal ganglion, VG: visceral ganglion, RPG: right parietal ganglion. Scale bar is 100µm. (n=12 individuals).

### Localization of innexin in the CNS

Because we found innexin paralogs could be expressed in the CNS, we sought to determine their localization at the cellular level. To this end, probes targeting unique regions in the *Lst Inx1* sequence were employed in ISH (Fig. 5) with sense probes used as a control (Fig. S3). ISH results showed differential localization of *Lst Inx1* within and between ganglia. For example, left and right pedal A (red), pedal E and F (yellow), and cerebral A cluster neurons (blue) have relatively high *Lst Inx1* localization, mostly concentrated on the plasma membrane. Interestingly, a higher expression of transcript is localized to the left pedal ganglia than the right pedal ganglia, indicating ganglionic heterogeneity in the expression of the same innexin paralog. Similar heterogenic membrane localization of *Lst Inx1* is also found in the electrically coupled cerebral A cluster.

Less membrane localization is found in the plasma membrane of visceral E, H, I, J, and K clusters, with the transcript being localized to the cytoplasm of some cells. Visceral L/M, right parietal B, and caudo-dorsal cluster neurons demonstrate intriguing results; these clusters strongly localize *Lst Inx1* but have not yet been revealed electrophysiologically to be coupled. It is interesting to note that the neurosecretory caudo-dorsal cells discharge at the same time when they release growth hormones during egg laying behavior (Kits and Bos, 1982; Roubos, 1976) suggesting a likely chance that they are electrically coupled. Electrophysiological experiments could support the hypothesis of electrical synapse abilities due to *Lst Inx1* localization.

Our results also demonstrate the lack of *Lst Inx1* localization on functionally defined electrical coupling cells. For example, right parietal dorsal 2 (RPD2) cell, located in the right parietal ganglion, is known to form very strong electrical synapses with ventral dorsal 1 (VD1) cell, located in the visceral ganglion (Soffe and Benjamin, 1980); however, neither cell localizes *Lst Inx1* transcript. Perhaps, then, a different innexin paralog is being localized to permit strong electrical coupling in these cells. These results support the exciting possibility of a connection between cell specificity in innexin paralog localization and cell-specific functions. Overall, localization of *Lst Inx1* was distributed throughout the entire *L. stagnalis* CNS. Because ganglia localized *Lst Inx1* in some cells but not others, further study of the remaining innexin paralogs could determine if the cells undetected by *Lst Inx1* probe would localize a different innexin transcript.

## Discussion

Gap junction-mediated electrical synapses in the nervous system are ubiquitous throughout vertebrates and invertebrates (Nagy et al., 2018; Stebbings et al., 2002). They play essential roles in development and complex behaviors in all animals including humans. Invertebrate models such as *L. stagnalis* contain large and functionally identified gap junction-forming neurons and can be used as valuable resources to explore gap junction formation and channel gating mechanisms. The current lack of molecular information on gap junctions in *L. stagnalis* as well as in many other invertebrate systems, however, severely prevents a comprehensive analysis of gap junction formation, transmission, and plasticity. To address this significant knowledge gap, we, for the first time, identified and characterized the expression of gap junction genes in *L. stagnalis*. It is our hope that this original molecular work will bring more research avenues to the gap junction field using the robust model *L. stagnalis* for comparative physiology, fundamental neurobiology, and biomedical research. To this end, we identified eight innexin paralogs by initial sequencing and BLAST analysis against the *L. stagnalis* genome. The innexins showed similarity with other invertebrate innexins and exhibited the same topology as invertebrate innexins, vertebrate connexins, and pannexins (the vertebrate homologues of innexins) (Baranova et al., 2004). Using RT-PCR and ISH, we provided evidence that innexin expression is paralog- and ganglia-specific, opening a potential link between innexin paralog expression and functional outcomes.

### Innexins in invertebrates

Our study revealed at least eight innexin paralogs present in *L. stagnalis*, which is fewer than 25 paralogs in *C. elegans* (Altun et al., 2009) and 21 paralogs in *H. verbana* (Kandarian et al., 2012) but similar to eight paralogs in *D. melanogaster* (Stebbings et al., 2002), eight paralogs in *O. bimaculoides* (Albertin et al., 2015), six paralogs in *C. borealis*, and six paralogs in the lobster *Homarus americanus* (Shruti et al., 2014). In vertebrates, multiple paralogous connexins and pannexins are also seen; the human genome contains 20 connexins, the mouse genome contains 19 connexins, and both genomes contain three known pannexins (Baranova et al., 2004; Eiberger et al., 2001; Yen and Saier, 2007). An interesting question remains, then, as to the evolutionary significance of the existence of various numbers of gap junction genes in different organisms. In addition to the well-accepted reasoning that one paralog can compensate for dysfunction of another paralog during gene loss or mutation-induced loss-of-function, evidence suggests the variation in connexin or pannexin gene number in vertebrates contributes to formation of heterotypic channels, leading to diverse channel functions, permeabilities, and gating mechanisms (Bukauskas and Verselis, 2004; Rackauskas et al., 2007). However, the physiological characteristics of diverse subunit combinations have yet to be fully explored in our and other invertebrate models.

A multiple sequence alignment revealed trends within the eight *L. stagnalis* paralog sequences. For example, a proline residue consistently located in the second transmembrane domain of our *L. stagnalis* innexins correlates to a proline found in the same domain in connexins. In connexins, this proline may be involved in voltage gating-associated conformational changes (Sansom and Weinstein, 2000), an idea that has yet to be fully studied in invertebrates. Some obvious differences between *L. stagnalis* innexins were present at the amino- and carboxyl-termini, a common theme among gap junction sequences (Bauer et al., 2005). Structural work in *C. elegans* by Oshima has suggested the involvement of the amino-terminus in the regulation of gap junction channel activity (Oshima et al., 2016). In addition, the carboxyl-terminus is thought to determine the functional variability in connexins, as it is the site for modification via phosphorylation (Giepmans, 2004). As such, the differences seen in the amino- and carboxyl-termini of our *L. stagnalis* sequences suggest potential differences in functionality of the gap junction protein channels formed. Because *L. stagnalis* neurons are large and cell culture of coupled neurons has been well established (Feng et al., 1997; Syed et al., 1990; Xu et al., 2014), an opportunity for future combination of molecular and electrophysiological analysis is possible.

The multiple sequence alignment also revealed that *Lst Inx7* is not as conserved as the other seven paralogs. At the serine/threonine amino acid site in the first transmembrane domain (big, green circle in Fig. 2B), *Lst Inx7* has a methionine. In the YY(x)W region in the second transmembrane domain, *Lst Inx7* has a phenylalanine instead of a tyrosine in the first position. These amino acid residues are two of the highly conserved residues in all invertebrate innexins (Phelan and Starich, 2001); therefore, the sequence differences between *Lst Inx7* and other innexins is unexpected, and it is not known if this counts for the minimal detection of its expression in tissues in our study (Fig. 4) as well as low read counts in RNA-sequencing data (Table S1). Nevertheless, it is interesting to note that one other innexin, *C. elegans Ce-inx-22*, also differs from typical innexin sequences at two amino acid residues: the first residue in the YY(x)W region in the second transmembrane domain and the proline position in the second extracellular loop (Phelan and Starich, 2001). In *C. elegans*, *Ce-inx-22* is expressed in germ cells, is thought to form heteromeric gap junctions with *Ce-inx-14*, and, along with *Ce-inx-14*, was screened as a negative regulator of oocyte maturation (Simonsen et al., 2014). An interesting future study could further explore the expression and role of *Lst Inx7* as a potential regulator of egg maturation to form a possible link between differences in amino acid residues and functional differences in paralogs.

### Evolutionary history of gap junctions in invertebrates

Our phylogenetic analysis demonstrated that all eight innexin genes in *L. stagnalis* were generated before its divergence from *A. californica*. *A. californica*, like *L. stagnalis*, is a gastropod but is a saltwater hare while *L. stagnalis* is a freshwater snail (Feng et al., 2009; Moroz et al., 2006). *L. stagnalis* and *A. californica* diverged approximately 237 million years ago (Hedges et al., 2015). With such a long period of time post-divergence, we must ask why both species retained multiple paralogs of innexins. It is tempting to assume that the paralogous innexins in *L. stagnalis* and *A. californica* have experienced functional diversifications, resulting in different roles in the development and function of electrical synapse, and thus have been retained during the evolution of *L. stagnalis* and *A. californica*.

It is also interesting that all invertebrate phyla examined form phylum-specific clades and exhibit similar gene duplication patterns of innexins. This suggests that gene duplication and functional differentiation are evolutionary driving forces and universal strategies for animal survival and function. Gene duplication is well accepted as a driving force of phenotypic evolution by generating raw genetic materials for functional innovation (Ohno, 1970). Functional innovation can be achieved by diversification of coding sequences and gene expression patterns (Zhang, 2003). The divergence of gene expression among innexin paralogous genes in *L. stagnalis* suggests that functional diversification occurred after the serial duplications of innexin genes during the evolution of *L. stagnalis*. The specific functions of each innexin gene in *L. stagnalis* still remain largely unclear. Future functional characterization of these innexin genes could provide further insight into understanding the evolution of gap junctions in invertebrates.

Another interesting trend emerges when comparing *O. bimaculoides* innexins to other Molluscan species. Five innexins in *O. bimaculoides* were duplicated, with no innexins in other molluscs being members of the same clade. In addition, clades were formed with *Lst Inx5, Lst Inx6, Lst Inx7*, and *Lst Inx8* in which no innexins from *O. bimaculoides* were members. Presumably, these results can be explained by two scenarios: *Lst Inx5, Lst Inx6, Lst Inx7*, and *Lst Inx8* and their related innexins were duplicated after the divergence of *O. bimaculoides* or *O. bimaculoides* lost the four related genes. Octopi are known for their highly intelligent behaviors and camouflage abilities (Gutnick and Kuba, 2018), and *O. bimaculoides* has been primarily studied for social behaviors that are very similar to humans. Results from these studies have demonstrated that the effect of serotonin on social behaviors and even the binding site in serotonin transporters is evolutionarily conserved between humans and *O. bimaculoides*; these same results have yet to be found in other molluscs (Edsinger and Dolen, 2018).

Perhaps, then, other highly intelligent characteristics seen in octopi can be attributed to differences in genetic makeup, like the additional five innexins genes in *O. bimaculoides* not related to other molluscan species. Incorporating innexin in other species of octopus in future phylogenetic analyses would be interesting to determine if the additional five innexin duplications are conserved across all octopus species, indicating a possible connection between behavior differences in the Mollusca phylum and genetics. In addition, while the evolutionary emergence of innexins, connexins, and pannexins has been studied, the functional abilities of each channel are still unclear. For example, innexins in the leech *H. medicinalis* can form gap junction channels, similar to connexins, as well as non-junctional channels, similar to pannexins (Bao et al., 2007). A phylogenetic analysis comparing innexins, connexins, and pannexins by functional ability could help explain why the three channel proteins have different functions.

### Paralog- and ganglion-specific expression and function

We wanted to delve further into the potential functional differences conferred by each *L. stagnalis* innexin paralog by first determining the paralog expression throughout tissue types. Our results demonstrated that all paralogs are expressed in every tissue but to different extents. In some cases, such as *Lst Inx4*, expression was highest in a single organ, the CNS. For *Lst Inx6*, though, the CNS had the least expression relative to other organs. This finding is not surprising considering the distribution pattern of gap junction genes in other invertebrates and vertebrates. For example, of the 21 innexins in *H. verbana*, only 11 are detectably expressed in the embryo CNS while five are expressed in the nephridia (Kandarian et al., 2012). The same variable gap junction gene distribution is found in human and mouse tissue (Oyamada et al., 2005).

Changes in gap junction expression are required for proper development, synaptic connections, and plasticity in invertebrates and vertebrates (Bhattacharya et al., 2019; Hall, 2017; Oyamada et al., 2005; Stebbings et al., 2002), and the differences in innexin expression levels found in *L. stagnalis* are likely required for proper organismal function under intrinsic, developmental and extrinsic, environmental regulations. Interestingly, *Lst Inx7* was downregulated compared to the other innexins in most tissues. This is consistent with the quantification analysis of previously published RNA-seq data showing no reads mapped to *Lst Inx7* (Table S1). However, it is important to note that this low expression shown in our study cannot exclude its expression and importance in other organs or under regulations by both developmental and environmental factors.

The effect of intrinsic and extrinsic factors on gap junction gene expression is an understudied field in invertebrates. Therefore, we wonder if other external or internal factors can help explain the differences seen in tissues of different snails. For example, hunger is known to change electrical synapse activity in *L. stagnalis*. Dyakonova et al. found that the firing rate of electrical synapse-forming pedal A neurons significantly increased when *L. stagnalis* were deprived of food for 42 hours (Dyakonova et al., 2015). However, to our knowledge, no studies have looked at the effect of hunger on gene expression; a slight possibility exists that the number of days since the snails’ last feeding could affect innexin expression. It would be very intriguing to test this postulation in future studies. In addition, an analysis of neuron-specific gene expression was recently completed in *C. elegans* which showed changes in innexin expression in response to transition to the dauer stage (Bhattacharya et al., 2019). Similar to *C. elegans*, knowledge of the expression patterns of innexin in *L. stagnalis* can be used to make strides in the understanding of how the electrical connectome is established throughout development and changes in response to external cues.

Finally, because we found evidence of potential paralog-specific functions at the tissue level, we wanted to establish if any differences also existed at the single cell level. Because our RNA-seq analysis of transcription abundance of innexin genes showed *Lst Inx1* to be the most highly expressed innexin (Table S1), we next performed ISH with DIG-labeled probes designed to target *Lst Inx1* transcript. *L. stagnalis* CNS consists of relatively large neurons with well-studied neural networks. This was a significant advantage to our ISH study because we knew the ganglionic location and function of neurons with the ability to form electrical synapses. As mentioned previously, left and right pedal A cluster neurons are electrically coupled cells involved in pedal cilia used for locomotion (Kyriakides et al., 1989), and accordingly, our ISH data localized *Lst Inx1* to the plasma membrane. We found ganglionic heterogeneity in these neurons as well as in cerebral A cluster neurons. In *A. californica*, intrinsic properties of individual neurons explained asymmetrical electrical coupling between neurons of the feeding motor network (Sasaki et al., 2013). In addition, in *L. stagnalis*, pedal A and cerebral A clusters form electrical synapses with cells in both their ipsilateral and contralateral counterparts (Kyriakides et al., 1989; Syed and Winlow, 1991). With these results in mind, the heteroganglionic localization of *Lst Inx1* could help explain the selective electrical connectome established in the pedal and cerebral ganglia. Likely, protein transcribed by *Lst Inx1* takes part in the formation of gap junctions in these ganglia; an antibody recognizing Lst Inx1 protein has the potential to confirm this hypothesis. If proven true, pedal A and cerebral A cells as well as other cells identified by this study to have *Lst Inx1* transcript localization could be used in further study of the voltage gating, permeability, and functional properties of the Lst Inx1 protein.

Localization of the innexin gene and protein paralogs are known to vary at the cellular level in invertebrates. In the CNS of *H. verbana*, innexins are expressed in a select number of cells. When ectopically expressed, though, *Hve-inx6* and *Hve-inx2* can form an electrical connection with cells to which they are not normally coupled. From these findings, the authors propose the expression of a specific innexin paralog is sufficient for electrical coupling (Firme et al., 2012). In *C. elegans*, nearly every cell type expresses at least one paralog of innexin, allowing the formation of heterotypic and heteromeric gap junctions (Hall, 2017). Our results further support the theme of cell-specific innexin paralog expression and localization which has potential for proper channel formation and changes in synaptic connection, ultimately leading to functional specificity of paralogs. Using the large, culturable neurons of our *L. stagnalis* model (Xu et al., 2014) and the cell-specific innexin expression data we present here, pathways, mechanisms, and factors regulating electrical synapse formation and function can be studied in a novel light: the role of innexin genes in electrosynaptogenesis.

## Conclusion

Although electrical synapses were discovered in 1959, very limited information is available on the genetic aspects of gap junction formation and plasticity (Furshpan and Potter, 1959); this is mostly due to the extreme complexity of nervous systems in most model organisms. Discoveries made in animals with simpler nervous systems are, therefore, profoundly critical for the understanding of gap junctions in human and vertebrates. Importantly, recent studies have found that certain interactions and pathways involving innexins are conserved among vertebrates (Alev et al., 2008; Bauer et al., 2006; Welzel and Schuster, 2018; Xu et al., 2001). Considering the prevalent roles of electrical networks in nervous system development and function, characterizing the molecular underpinning of electrical synapses is urgently warranted. If detailed genetic information is available, powerful genetic tools allow an in-depth analysis and fine dissection of cellular pathways for understanding the basic mechanisms of physiology and pathology of gap junctions in our and other model organisms. Compared to the established models of *D. melanogaster* and *C. elegans*, *L stagnalis’* large neurons, functionally defined network, and simple behaviors, together with its powerful synapse culture model and electrophysiology assays, provide a unique opportunity to study the molecular, synaptic, and physiological mechanisms related to learning and memory as well as neurobiological diseases. The availability of *L. stagnalis* innexins provided by the present study will aid our ability to study the molecular mechanisms related to gap junction formation and functions and eventually decipher their contribution to physiology and pathophysiology of the nervous system. With our *L. stagnalis* model, we now have the means to answer specific questions such as “What other transcription factors or proteins are used to regulate innexin expression and gap junction formation?”, “Is there a compensatory mechanism used when one innexin paralog is inhibited, such as in a diseased state, to allow normal functioning?”, and “Are these mechanisms and pathways conserved in vertebrate animals and humans?” Answers to these questions are critical to our understanding of the expression and function of gap junction genes and proteins, as well as inferring their evolutionary history and functional diversification in animals.

## Acknowledgments

The authors would like to thank Dr. Frank Visser for intellectual help with designing and troubleshooting 3’ and 5’ RACE primers.

## Data Availability

The genes identified in this study are available in GenBank under accession numbers outlined in Table 1.

---

**Supplementary Table 1.** Transcription level of innexin genes of the CNS in *L. stagnalis* based on RNA-sequencing data

| **Gene** | **Transcript Length** | **Mapped Read Count** | **RPKM** |
| --- | --- | --- | --- |
| *Lst Inx1* | 1134 | 698 | 7.52 |
| *Lst Inx2* | 1065 | 11 | 0.13 |
| *Lst Inx3* | 921 | 108 | 1.43 |
| *Lst Inx4* | 855 | 188 | 2.68 |
| *Lst Inx5* | 990 | 131 | 1.62 |
| *Lst Inx6* | 1101 | 1 | 0.01 |
| *Lst Inx7* | 678 | 0 | 0 |
| *Lst Inx8* | 1209 | 0 | 0 |

**Supplementary Figure 1.**
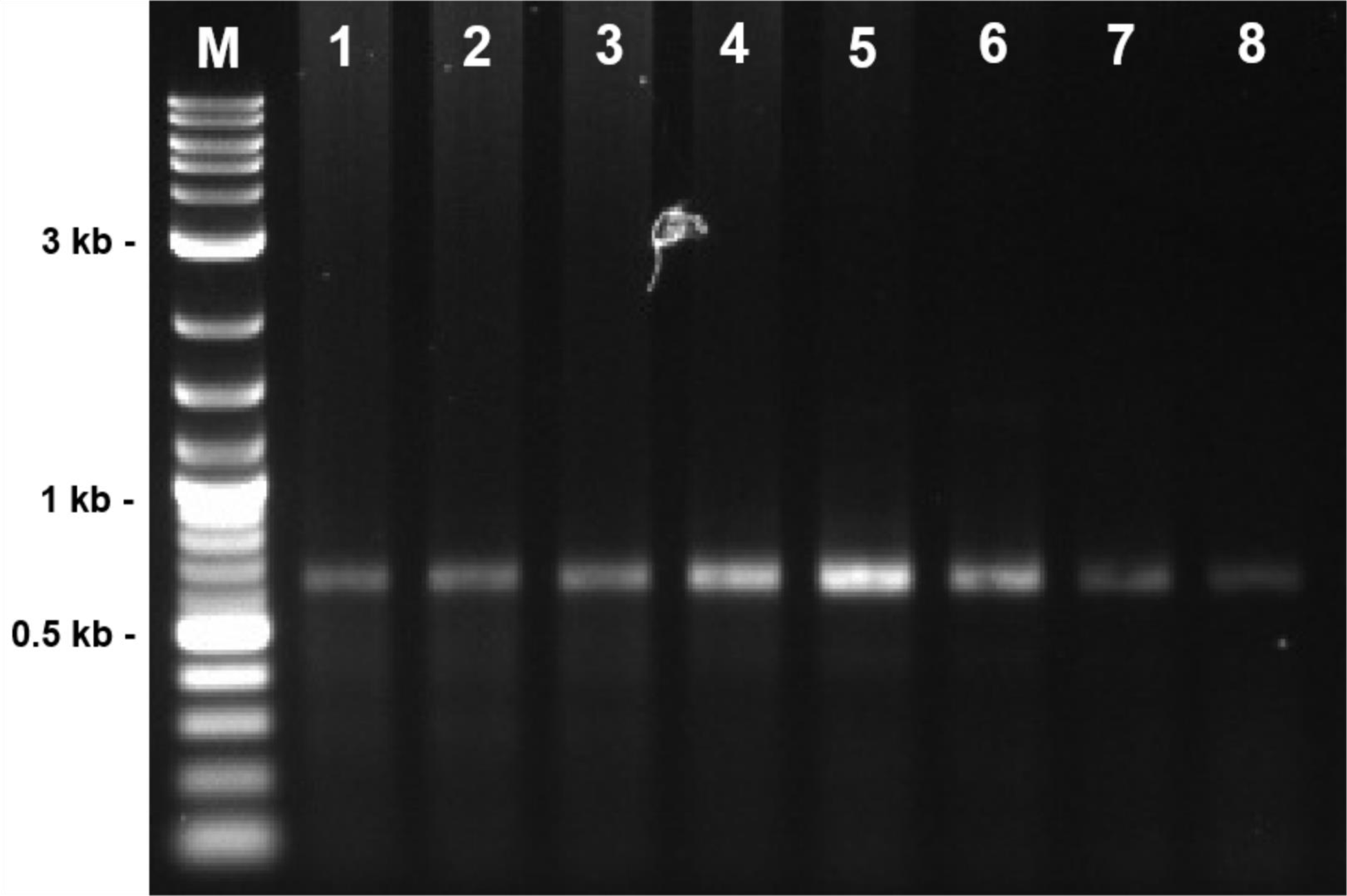
PCR with degenerate primers created for *C. borealis* revealed a partial sequence of innexin in *L. stagnalis*. The CNS of 12 snails was dissected, and RNA was extracted. Annealing temperatures in PCR varied in lanes one through eight, but all settings resulted in the same-sized partial sequence: lane 1: 40.0°C; lane 2: 40.2 °C; lane 3: 41.3°C; lane 4: 43.1°C; lane 5: 45.4°C; lane 6: 48.0°C; lane 7: 50.7°C; lane 8: 53.5°C. M=molecular ladder.

**Supplementary Figure 2.**
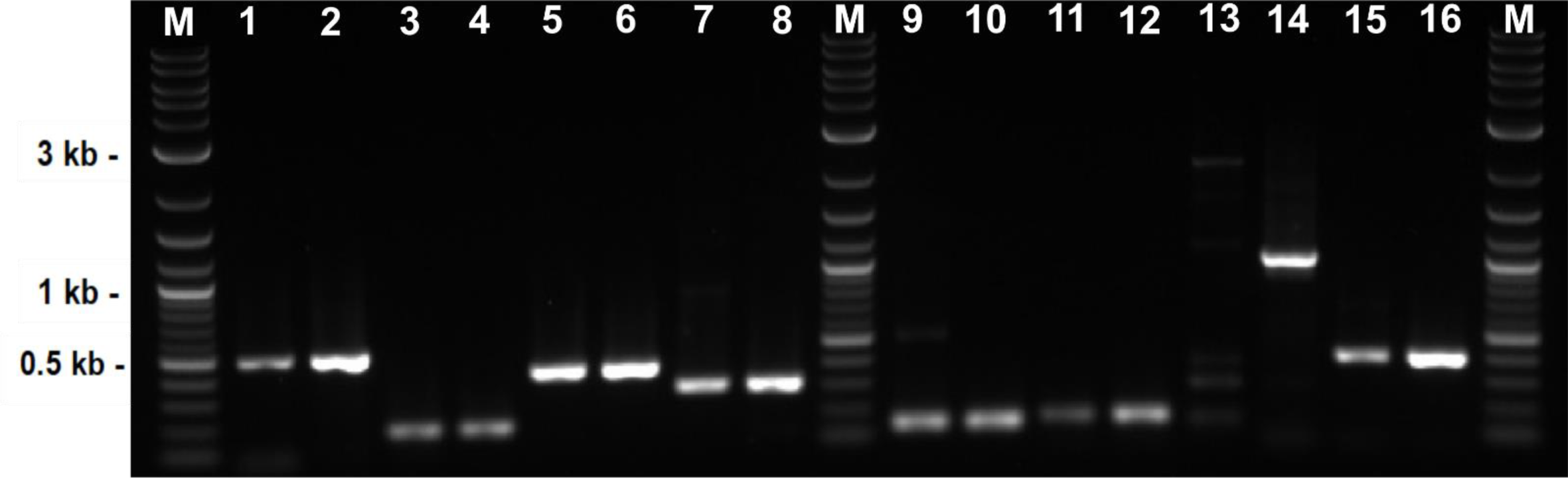
Representative agarose gel shows the results of RT-PCR RNA and gDNA reactions. Here, RNA and gDNA was extracted from *L. stagnalis* CNS, and primers specific to each paralog were used (Table 4). Lane 1: *Lst Inx1* RNA; lane 2: *Lst Inx1* gDNA; lane 3: *Lst Inx2* RNA; lane 4: *Lst Inx2* gDNA; lane 5: *Lst Inx3* RNA; lane 6: *Lst Inx3* gDNA; lane 7: *Lst Inx4* RNA; lane 8: *Lst Inx4* gDNA; Lane 9: *Lst Inx5* RNA; lane 10: *Lst Inx5* gDNA; lane 11: *Lst Inx6* RNA; lane 12: *Lst Inx6* gDNA; lane 13: *Lst Inx7* RNA; lane 14: *Lst Inx7* gDNA; lane 15: *Lst Inx8* RNA; lane 16: *Lst Inx8* gDNA. Of note, the primer pair used for *Lst Inx7* is intron-spanning, and the resulting band includes one 558bp intron that is spliced out during mRNA production. Therefore, the *Lst Inx7* band in gDNA reactions is 1011bp rather than 446bp, like in RNA reactions. Designing an intron spanning set of primers added a level of control to the highly downregulated *Lst Inx7* to ensure the amplified RNA did not include any gDNA. The RNA and gDNA band intensities were used to create Fig. 4. M=molecular ladder.

**Supplementary Figure 3.**
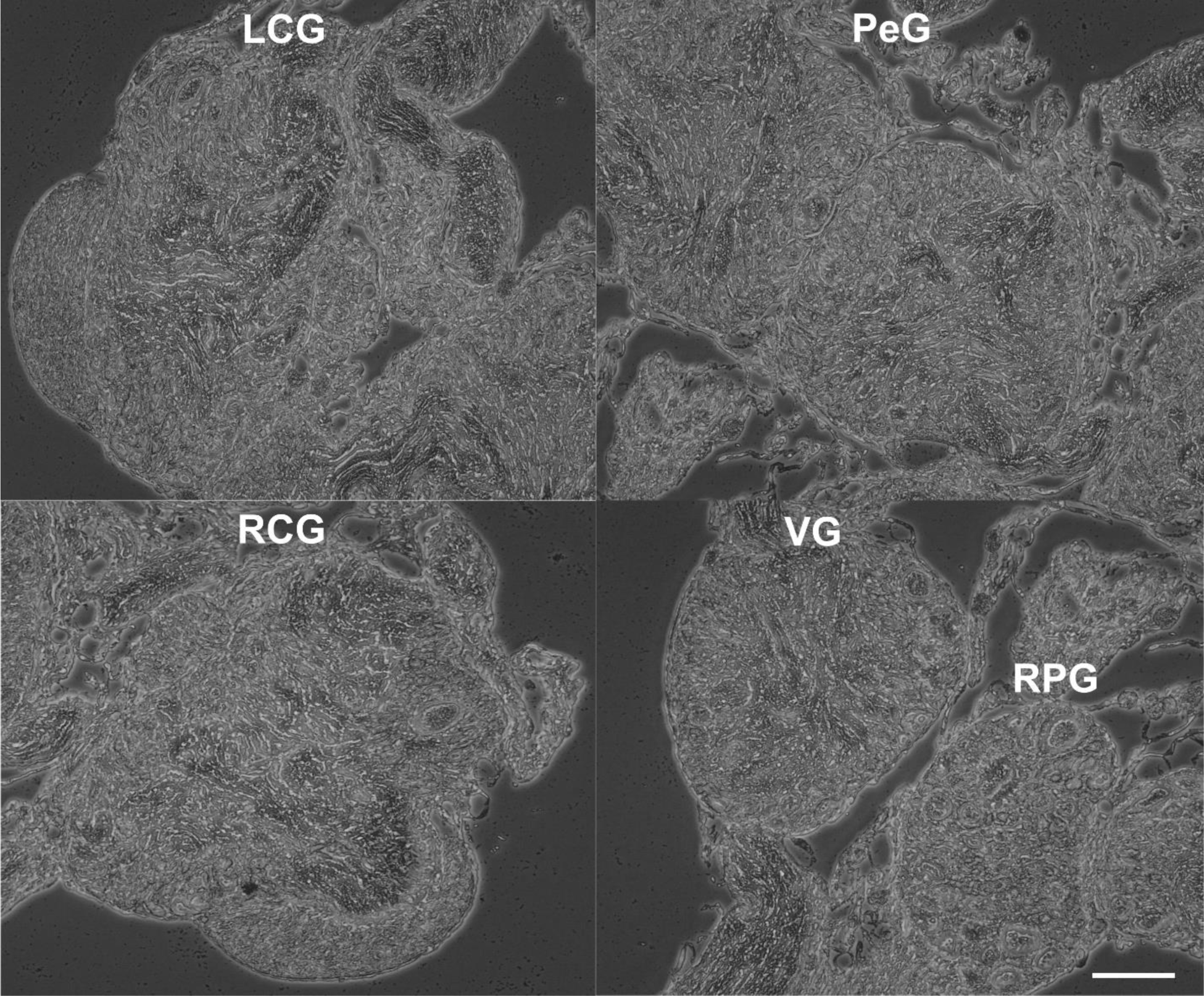
Sense probes were used as controls for *in situ* hybridization. The same ganglia identified in Fig. 5 are shown here. LCG: left cerebral ganglion, RCG: right cerebral ganglion, PeG: pedal ganglion, VG: visceral ganglion, RPG: right parietal ganglion. Scale bar is 100µm. (n=12 individuals)

